# Pelvic Ring Fractures: Does Spinopelvic Alignment Affect Sacral and Lumbopelvic Fixation Stability?

**DOI:** 10.1101/2024.10.17.618950

**Authors:** Sudharshan Tripathi, Yogesh Kumaran, Sophia M. Soehnlen, Amey Kelkar, Toshihiro Seki, Takashi Sakai, Vijay Goel, Carmen E. Quatman, Norihiro Nishida

## Abstract

**Background Context:** Pelvic ring is a complex boney and ligamentous complex and its structural integrity is paramount to be able to withstand the high forces. Pelvic ring fractures, which can result from low-energy impacts or high-velocity injuries and falls from great heights, can severely impair the structural integrity of the pelvic ring. These injuries can be fatal, especially in geriatric patients, with mortality rates ranging between 10% and 16%. Pelvic ring injuries require immediate stabilization, typically performed through internal fixation, to maintain circulatory balance and achieve anatomical restoration of the pelvis.

**Methods:** A previously validated FE model of the spine, pelvis, and femur model was used to evaluate how different fixation techniques for pelvic ring injury affect spinopelvic biomechanics. The SS, PI, PT and LL of the normal, intact model was modified to generate two more SS models for this study. One modified model with an SS of 20º was considered the low SS model (SSD), while the second modified model with an SS of 32º was termed the high SS model (SSI). A unilateral pelvic ring fracture was simulated by resecting the left side of the sacrum and pelvis. The fractures were stabilized using two different posterior stabilization techniques 1) The fractures were stabilized using two different posterior stabilization techniques, and 2) L5-Ilium posterior screw fixation without cross connector (L5_PF_WO_CC).

**Results:** SSI demonstrated the higher range of motion, followed by Normal model and least by SSD model at right Sacroiliac Joint. No differences were seen between with and without cross connector model for range of motion at right SIJ. For L5-S1 motion models treated with cross connectors demonstrated the least motion compared to those treated without cross connectors for all configurations and loading conditions. For the sacrum fracture, models with high sacral slopes recorded the least horizontal displacement, followed by decreased sacral slope models for both configurations and loading conditions.

## Introduction

The spine, pelvis, and hip form an interconnected mechanism essential for maintaining biomechanical standing balance in humans. This balance is largely attributed to the mechanical stability offered by the pelvic ring, Pelvic ring is a complex boney and ligamentous complex and its structural integrity is paramount to be able to withstand the high forces[1]. Pelvic ring fractures, which can result from low-energy impacts or high-velocity injuries and falls from great heights, can severely impair the structural integrity of the pelvic ring [1, 2]. These injuries can be fatal, especially in geriatric patients, with mortality rates ranging between 10% and 16%[3]. Pelvic ring injuries require immediate stabilization, typically performed through internal fixation, to maintain circulatory balance and achieve anatomical restoration of the pelvis [4]. Several techniques for pelvic ring fracture fixation have been used with good results, including trans-iliac–trans-sacral screws, ilio-sacral (IS) screws, trans iliac rod and screw fixation (TIF), and lumbopelvic fixation (LP). [5-7] [7] [8, 9] [10] However, the spinopelvic anatomy differs significantly from patient to patient and drives the choice of fixation technique which may be used for a particular anatomy.. Usually in treatment of pelvic fractures, fixation at the fracture site and the reduction of the fracture site are most important factors when the surgical pre-planning is performed. Key spinopelvic parameters such as Lumbar lordosis (LL), pelvic incidence (PI), pelvic tilt (PT), and sacral slope (SS) are crucial in defining sagittal balance of the spine[11, 12]. Even though previous research has shown that varying SS and PT angles alter the biomechanics of the lumbar spine, and sacroiliac joint (SIJ) [13], currently pre-existing spinopelvic parameters are given less-of-an importance in surgical pre-planning for the treatment of pelvic ring fractures even though they may alter the mechanical alignment and function post-surgery.

To our knowledge, no studies have analyzed the impact on the mechanics of the SIJ, and intervertebral disc (IVD) after fracture fixation treatments for unstable type IIC pelvic fractures with varying SS and PT angles. We hypothesize that sagittal balance might also influence pelvic fracture fixation mechanics. Additionally, we hypothesize that each fixation technique may alter the biomechanics of surrounding joint structures such as the intervertebral discs, hip joint differently and may lead to differing long term degenerative effects on these surrounding joints.

We propose the use of a finite element (FE) lumbar spinopelvic model with an unstable type IIC pelvic fracture, combined with varying SS angles and multiple motions, to study the effects on biomechanics of the IVD, and SIJ, joints after fracture fixation treatments for unstable type IIC pelvic fractures [2].

## Materials and Methods

A previously validated FE model of the spine, pelvis, and femur model [13]was used to evaluate how different fixation techniques for pelvic ring injury affect spinopelvic biomechanics. The model was developed from a computed tomography (CT) scan of a healthy patient without any abnormalities or bone deformities. MIMICS software (Materialise, Leuven, Belgium) was used to generate the 3D geometry of the spine and pelvis. Geomagic Studio software (Raindrop Geomagic Inc., Raleigh, North Carolina, USA) was employed to smooth the geometry. The smoothed intervertebral discs were meshed using IAFEMESH software (University of Iowa, Iowa City, Iowa, USA), while the vertebrae and pelvis were meshed using Hypermesh (Altair Engineering Inc., Troy, Michigan, USA). A 1 mm thick mesh of cortical bone was assigned to the vertebrae and pelvis which surrounded the cancellous bone. The meshed model of the spine and pelvis was assembled using Abaqus FEA software (Dassault Systemes/Simulia Inc., Providence, Rhode Island, USA). Linear hexahedral elements were assigned for the cortical bone of the vertebrae and intervertebral discs, while tetrahedral elements were assigned for the cancellous bone of the vertebrae and pelvis.

Ligamentous tissues, including the anterior longitudinal ligament, posterior longitudinal ligament, interspinous ligament, supraspinous ligament, capsular ligament, and ligamentum flavum, were modeled as truss elements. Similarly, the ligaments of the sacroiliac joint (SIJ)—namely, the anterior sacroiliac ligament, sacrospinous ligament, interosseous ligament, posterior sacroiliac ligament, short posterior ligament, and sacrotuberous ligament—were also modeled as truss elements. Nonlinear soft contact interactions were defined for the sacroiliac joints, spine facets, articular cartilages, and pubic symphysis.

The femur was positioned relative to pelvis according to a study conducted by Wu et al. [14] by aligning the center of rotation between femoral head and pelvis. The hip joint was validated with data obtained from Anderson et. al. [15] by comparing the FE results of hip contact stress, and area for walking, ascending stairs, and descending stairs with their results. This model was considered as the normal, intact model. The spinopelvic parameters of this model were SS= 26º LL= 42º, PI= 37º, and PT= 11º respectively [13].

**Figure 1.**
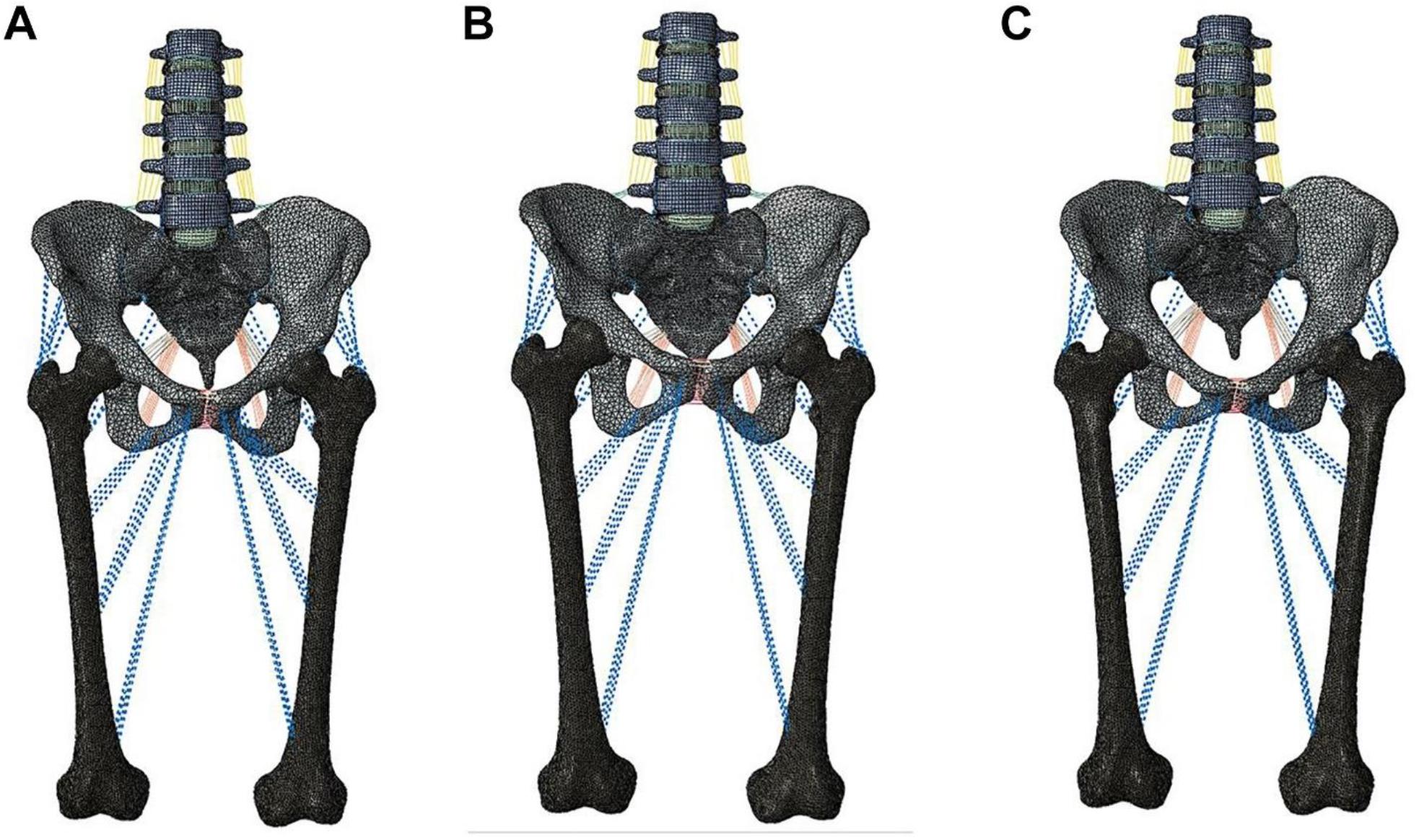
Intact model developed from CT images with no abnormalities (a)indicating Intact model, (b) indicating low sacral slope, and (c) indicating high sacral slope.

### Material Properties

The material properties of the FE model are summarized in Table [1]. All the material properties used for the FE model were obtained from previous literature [16-22].

**Table 1.** Finite Element Model Material Properties.

| Component | Material Properties | Constitute Relation | Element Type |
| --- | --- | --- | --- |
| Vertebral Cortical Bone | $E = 12,000 \text{ MPa}$<br>$\nu = 0.3$ | Isotropic, Elastic | 8 Node Brick Element (C3D8) |
| Vertebral Cancellous Bone | $E = 100 \text{ MPa}$<br>$\nu = 0.2$ | Isotropic, Elastic | 4 Node Tetrahedral Element (C3D4) |
| Pelvis Cortical Bone (Sacrum, Ilium) | $E = 17,000 \text{ MPa}$<br>$\nu = 0.3$ | Isotropic, Elastic | 4 Node Tetrahedral Element (C3D4) |
| Sacrum Cancellous Bone | Heterogenous | Isotropic, Elastic | 4 Node Tetrahedral Element (C3D4) |
| Ilium Cancellous Bone | E = 70 MPa | Isotropic, Elastic |  |
| | $\nu = 0.2$ | | 4 Node Tetrahedral Element (C3D4) |
| Femur Cortical Bone | E = 17,000 MPa | Isotropic, Elastic | 4 Node Tetrahedral Element (C3D4) |
| | $\nu = 0.29$ | | |
| Femur Cancellous Bone | E = 100 MPa | Isotropic, Elastic | 4 Node Tetrahedral Element (C3D4) |
| | $\nu = 0.2$ | | |
| Ground Substance of Annulus Fibrosis | $c_{10} = 0.035$ | | |
| | $k_1 = 0.296$ | Hyper elastic anisotropic (HGO) | 8 Node Brick Element (C3D8) |
| | $k_2 = 65$ | | |
| Nucleus Pulposus | E = 1 MPa | Isotropic, Elastic | 8 Node Brick Element (C3D8) |
| | $\nu = 0.499$ | | |
| Anterior Longitudinal | 7.8 MPa (<12%),<br>20 MPa (>12%) | Non-linear Hypo elastic | Truss Element (T3D2) |
| Posterior Longitudinal | 10 MPa (<11%),<br>20 MPa (>11%) | Non-linear Hypo elastic | Truss Element (T3D2) |
| Ligamentum Flavum | 15 MPa (<6.2%),<br>19.5 MPa (>6.2%) | Non-linear Hypo elastic | Truss Element (T3D2) |
| Intertransverse | 10 MPa (<18%),<br>58.7 MPa (>18%) | Non-linear Hypo elastic | Truss Element (T3D2) |
| Interspinous | 10 MPa (<14%),<br>11.6 MPa (>14%) | Non-linear Hypo elastic | Truss Element (T3D2) |
| Supraspinous | 8 MPa (<20%), 15 MPa (>20%) | Non-linear Hypo elastic | Truss Element (T3D2) |
| Capsular | 7.5 MPa (<25%),<br>32.9 MPa (>25%) | Non-linear Hypo elastic | Truss Element (T3D2) |
| Anterior SIJ | 125 MPa (5%),<br>325 MPa (>10%),<br>316 MPa (>15%) | Non-linear Hypo elastic | Truss Element (T3D2) |
| Short Posterior SI | 43 MPa (5%),<br>113 MPa (>10%),<br>110 MPa (>15%) | Non-linear Hypo elastic | Truss Element (T3D2) |
| Long Posterior SI | 150 MPa (5%),<br>391 MPa (>10%),<br>381 MPa (>15%) | Non-linear Hypo elastic | Truss Element (T3D2) |
| Interosseous | 40 MPa (5%), | Non-linear Hypo elastic | Truss Element (T3D2) |
|  | 105 MPa (>10%),<br>102 MPa (>15%) |  |  |
| Sacrospinous | 304 MPa (5%),<br>792 MPa (>10%),<br>771 MPa (>15%) | Non-linear Hypo elastic | Truss Element (T3D2) |
| Sacro tuberos Ligament | 326 MPa (5%),<br>848 MPa (>10%),<br>826 MPa (>15%) | Non-linear Hypo elastic | Truss Element (T3D2) |
| Gluteus Maximus | k = 344 N/mm |  | Connector Element |
| Gluteus Medius | k = 779 N/mm |  | Connector Element |
| Gluteus Miniums | k = 660 N/mm |  | Connector Element |
| Psoas Major | k = 100 N/mm |  | Connector Element |
| Adductor Magnus | k = 257 N/mm |  | Connector Element |
| Adductor Longus | k = 134 N/mm |  | Connector Element |
| Adductor Brevis | k = 499 N/mm |  | Connector Element |
| Rods (Titanium) | E = 120,000 MPa | Isotropic, Elastic | Hexahedral Element |
| | $\nu = 0.3$ | | |
| Pedicule Screws (Titanium) | E = 120,000 MPa | Isotropic, Elastic | Hexahedral Element |
| | $\nu = 0.3$ | | |

### Models with Varying Sacral Slopes

The SS, PI, PT and LL of the normal, intact model was modified to generate two more SS models for this study. One modified model with an SS of 20º was considered the low SS model (SSD), while the second modified model with an SS of 32º was termed the high SS model (SSI). The spinopelvic parameters for the SSD model were SS = 20º, LL = 36º, PI = 36º, and PT = 17º, whereas for the SSI model, the parameters were SS = 32º, LL = 42º, PI = 37º, and PT = 5º.

**Figure 2.**
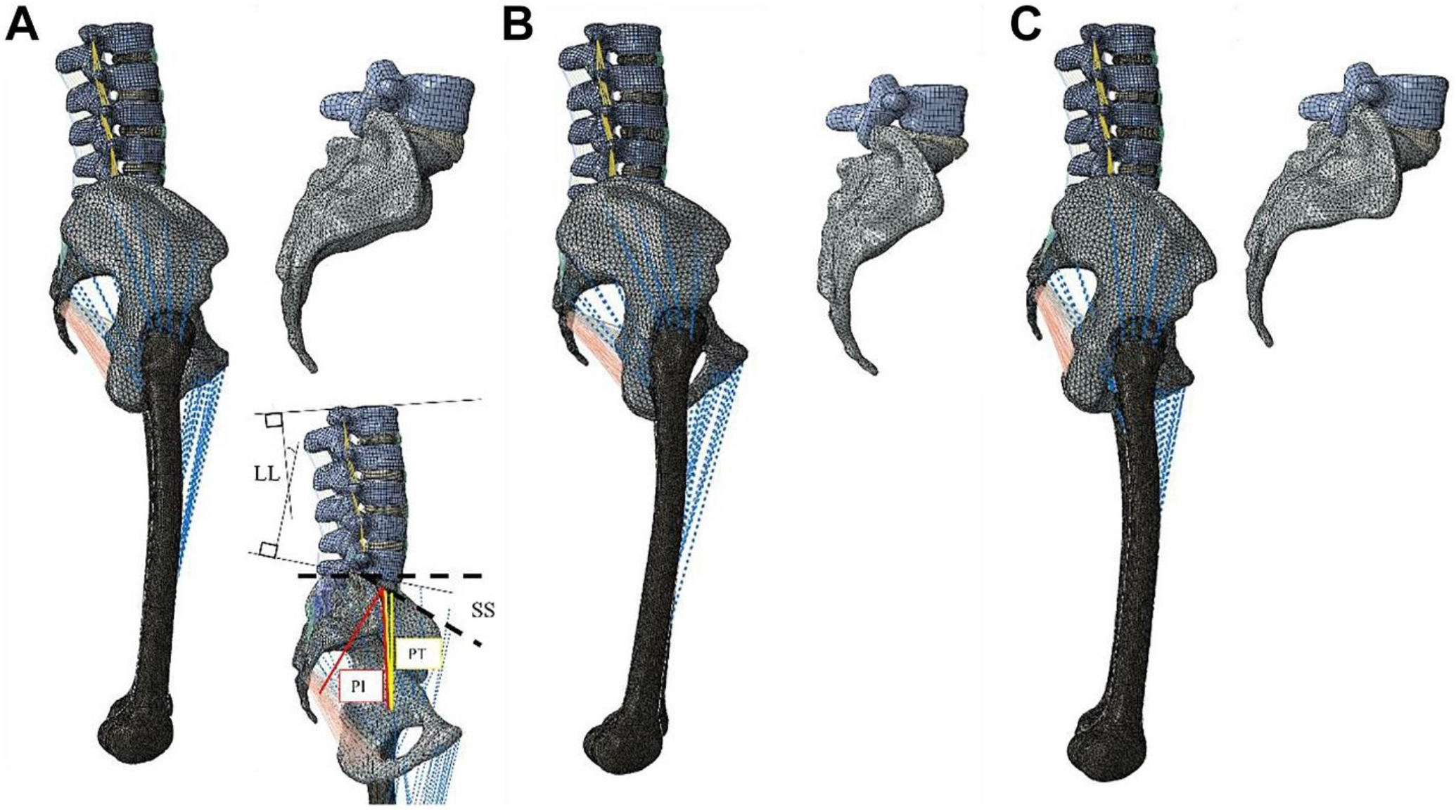
Lateral view of spine pelvis hip finite element mode (a) representing Intact model with sacral slope (SS) of 26 degrees, (b)representing low sacral slope model (SSD) with SS of 20 degrees, and (c) representing high sacral slope model (SSI) o sacral slope of 32 degrees.

### Simulation of Fracture

A unilateral pelvic ring fracture was simulated on the left side of the sacrum by resecting elements in the finite element (FE) model of the spine and pelvis for all cases with different SS. A type IIC fracture, as per the Rommens classification, was simulated with a gap of approximately 1.7 mm [2]. Surface-to-surface interaction with hard contact was defined between the fractured surfaces.

**Figure 3.**
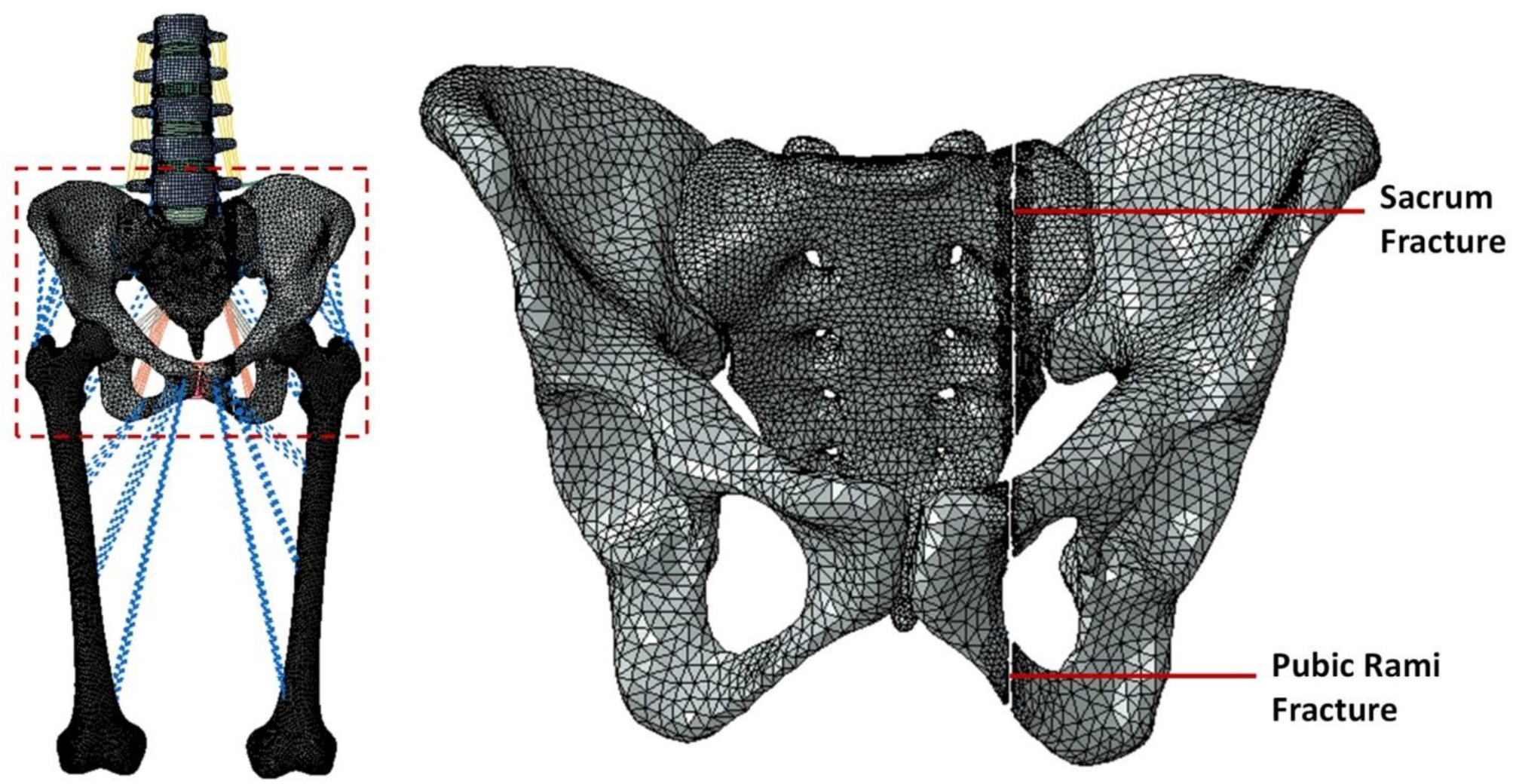
Simulation of the Rommens Type IIC Pelvic ring fracture on the left side.

### Simulation of Different Spinopelvic Fixations

The fractures were stabilized using two different posterior stabilization techniques utilizing the Galveston approach whereby transpedicular screws are implanted in the lower lumbar spine and supplemented with additional screws in the posterior ilium [23].

- L5-Ilium posterior screw fixation with cross connector (L5_PF_W_CC): Bilateral posterior screw fixation was performed from L5 to the ilium. The pedicle screws were connected to spinal rods, and a cross connector was placed at the S1 level to connect the two rods. In the results section, the fixation of the normal SS model with L5_PF_W_CC was termed as normal (L5_ILIUM_FIX_W_CC). For the SSD and SSI models, it was indicated as SSD (L5_ILIUM_FIX_W_CC) and SSI (L5_ILIUM_FIX_W_CC), respectively.
- L5-Ilium posterior screw fixation without cross connector (L5_PF_WO_CC): Bilateral posterior screw fixation was performed from L5 to the ilium. The pedicle screws were connected to spinal rods. In the results section, the fixation of the normal SS model with L5_PF_WO_CC was termed as normal (L5_ILIUM_FIX_WO_CC). For the SSD and SSI models, it was indicated as SSD (L5_ILIUM_FIX_WO_CC) and SSI (L5_ILIUM_FIX_WO_CC), respectively.

Screws, rods, and cross-connectors were assigned Ti-6Al-4V material properties. A rigid tie interaction was used to simulate the fixation between rods and screws, as well as between rods and cross-connectors. To simulate screw fixation in bone, a screw-bush and bush-bone interface was utilized. A tie constraint was applied between the outer surface of the screw and the inner surface of the bush, while a coupling constraint was used between the bush and the bone.

**Figure 4.**
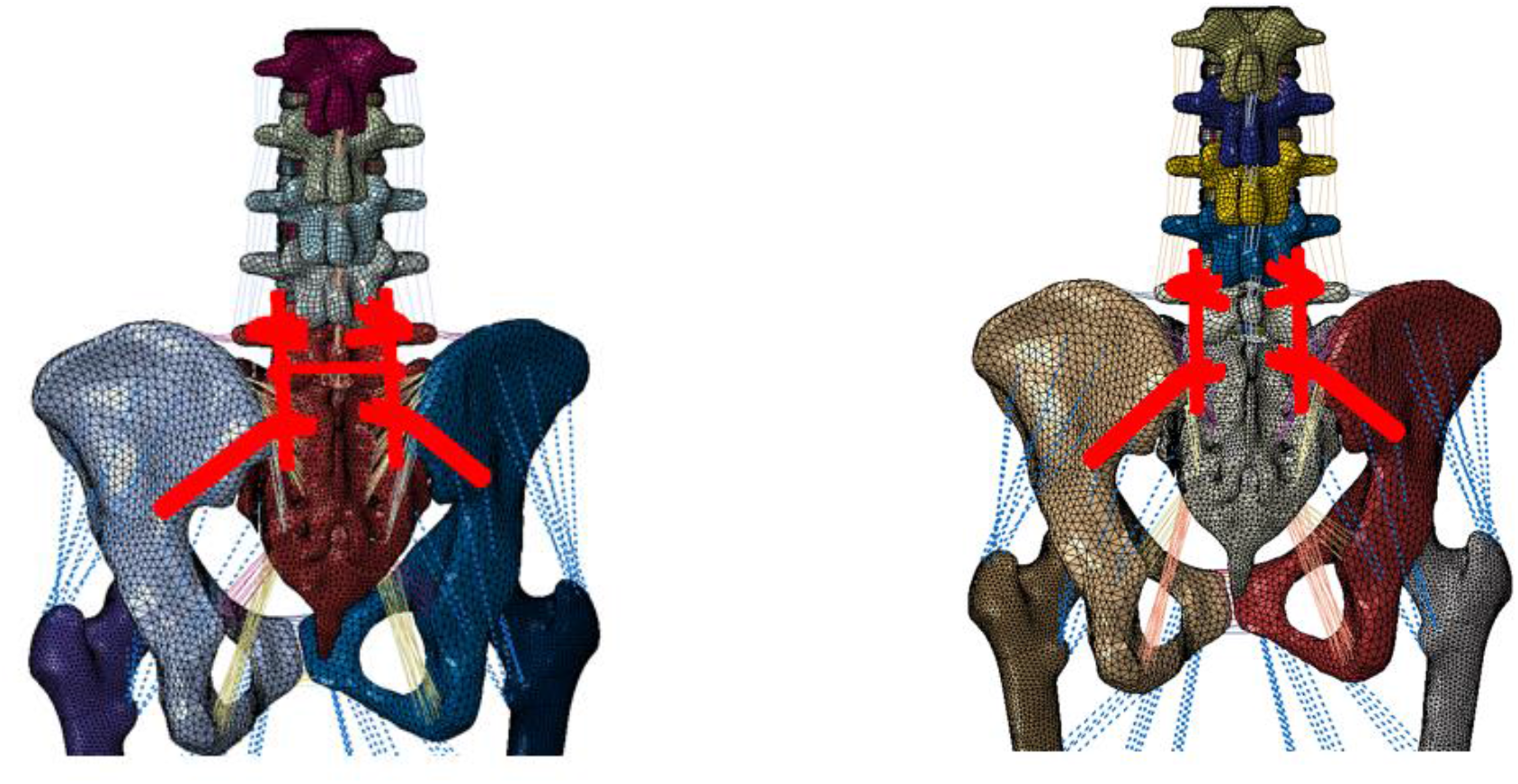
Stabilization of the Pelvic ring fracture with and without cross connector.

### Loading and Boundary Conditions

To replicate the weight of the upper trunk and muscle forces, a compressive follower load of 400N was applied to all models. Additionally, a bending moment of 7.5 Nm was applied to the superior surface of the L1 vertebra to simulate flexion/extension, lateral bending, and axial rotation. To simulate a two-leg stance posture, the distal portion of the femur was immobilized in all degrees of freedom [24].

## Data analysis

The stabilization offered by the 2 configurations for models with different SS was quantified by analyzing the horizontal and vertical displacement across the sacral and pelvic fractures. The contralateral side (right) SIJ range of motion (ROM), L5-S1 ROM, L5-S1 nucleus stress, SIJ stress, and stress on the rods was recorded and compared across two configurations for different SS models.

## Results

### Range of Motion

#### Right SIJ ROM (Figure 5)

The right SIJ ROM for all configurations and motions was less than 0.5 degrees. Flexion and extension showed the least motion (<0.25 degrees) for all configurations, while the greatest motion was seen in lateral bending (around 0.5 degrees). Under flexion, extension, and lateral bending, the decreased sacral slope model either fixed with cross connector or no cross connector showed the least motion compared to normal and the increased sacral slope models (< 0.25 degrees). For axial rotation, the increased sacral slope model treated with either cross connector or no cross connector resulted in the least motion (<0.1 degrees) compared with the normal (<0.25 degrees) and decreased sacral slope models (< 0.3 degrees). In lateral bending and axial rotation, models treated with cross connectors showed reduced motion compared to models without cross connectors for the normal and decreased sacral slope models. For flexion and extension, no minimal change in motion was observed between models treated with either a cross connector or no cross connector for all cases, i.e., normal, decreased, and increased sacral slope models. The left SIJ ROM was not recorded as the motion was relatively low due to the adjacent sacral fracture.

**Figure 5.**
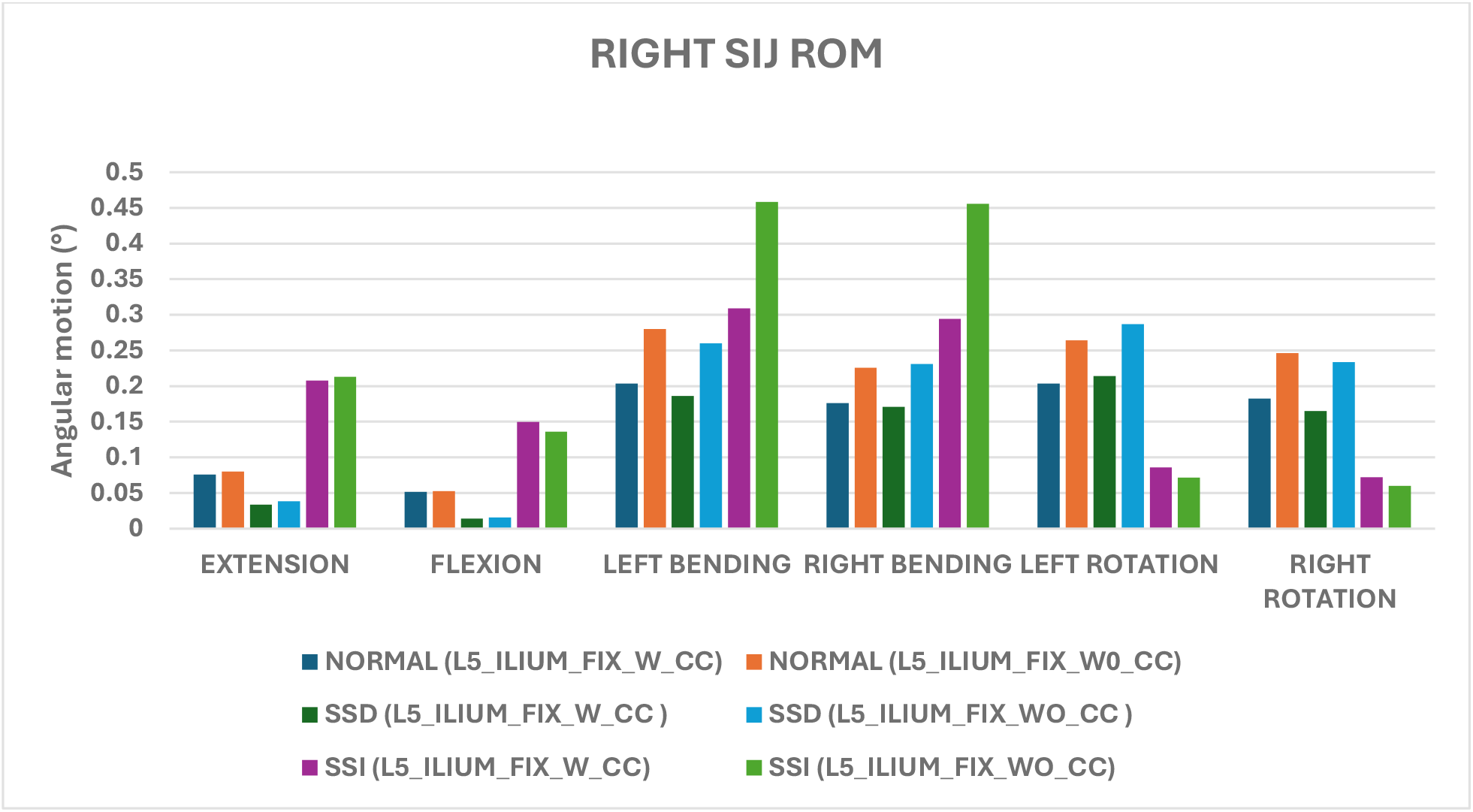
Comparison of ROM at right SIJ at 7.5Nm moment with 400N follower load under two leg stance condition for three different sacral slopes stabilized with and without cross-connector.

#### L5-S1 ROM (Figure 6)

The ROM of L5-S1 for all configurations was less than (<0.5 degrees) in all loading conditions. Lateral bending showed the least motion (<0.35 degrees) for all configurations, while axial rotation resulted in the largest motion (< 0.45 degrees). Motion for flexion and extension were similar for all configurations. Models treated with cross connectors demonstrated the least motion compared to those treated without cross connectors for all configurations and loading conditions.

**Figure 6.**
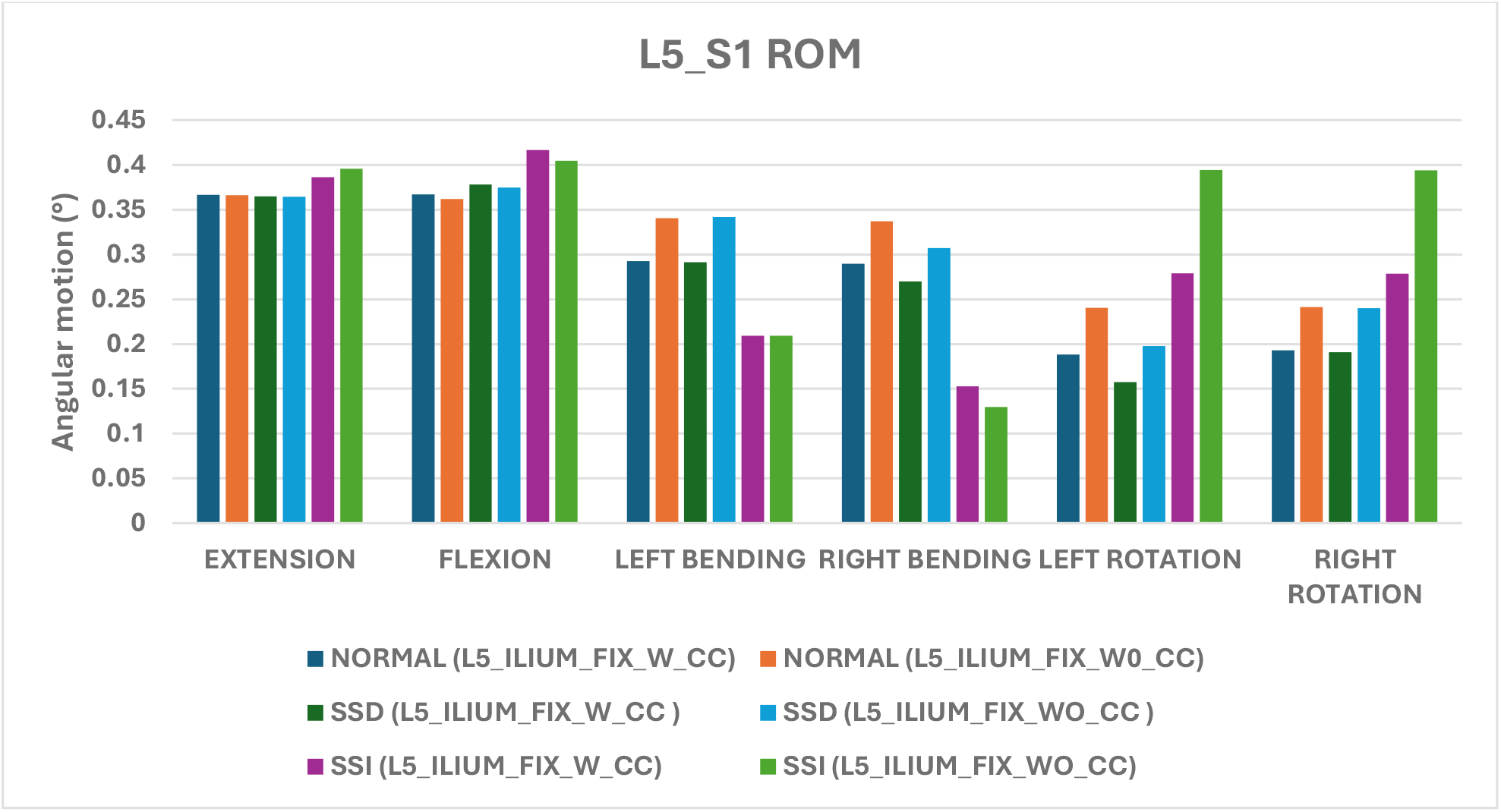
Comparison of ROM of L5-S1 at 7.5Nm moment with 400N follower load under two leg stance condition for three different sacral slopes stabilized with and without cross-connector.

Extension/flexion and axial rotation models with high sacral slope resulted in the greatest motion than models with normal and decreased sacral slopes treated with either a cross connector or no cross connector.

#### L5-S1 Nucleus Stress (Figure 7)

Normal sacral slope models showed the smallest L5-S1 nucleus stress (<1 MPa) for both configurations in all the loading conditions, followed by the low sacral slope models (< 2 MPa) in all loading conditions and both fixation configurations. The high sacral slope models resulted in larger stress (< 3 MPa) for both configurations and all loading conditions. Models with cross connectors resulted in less nucleus stress than those without cross connectors for all loading conditions and cases.

**Figure 7.**
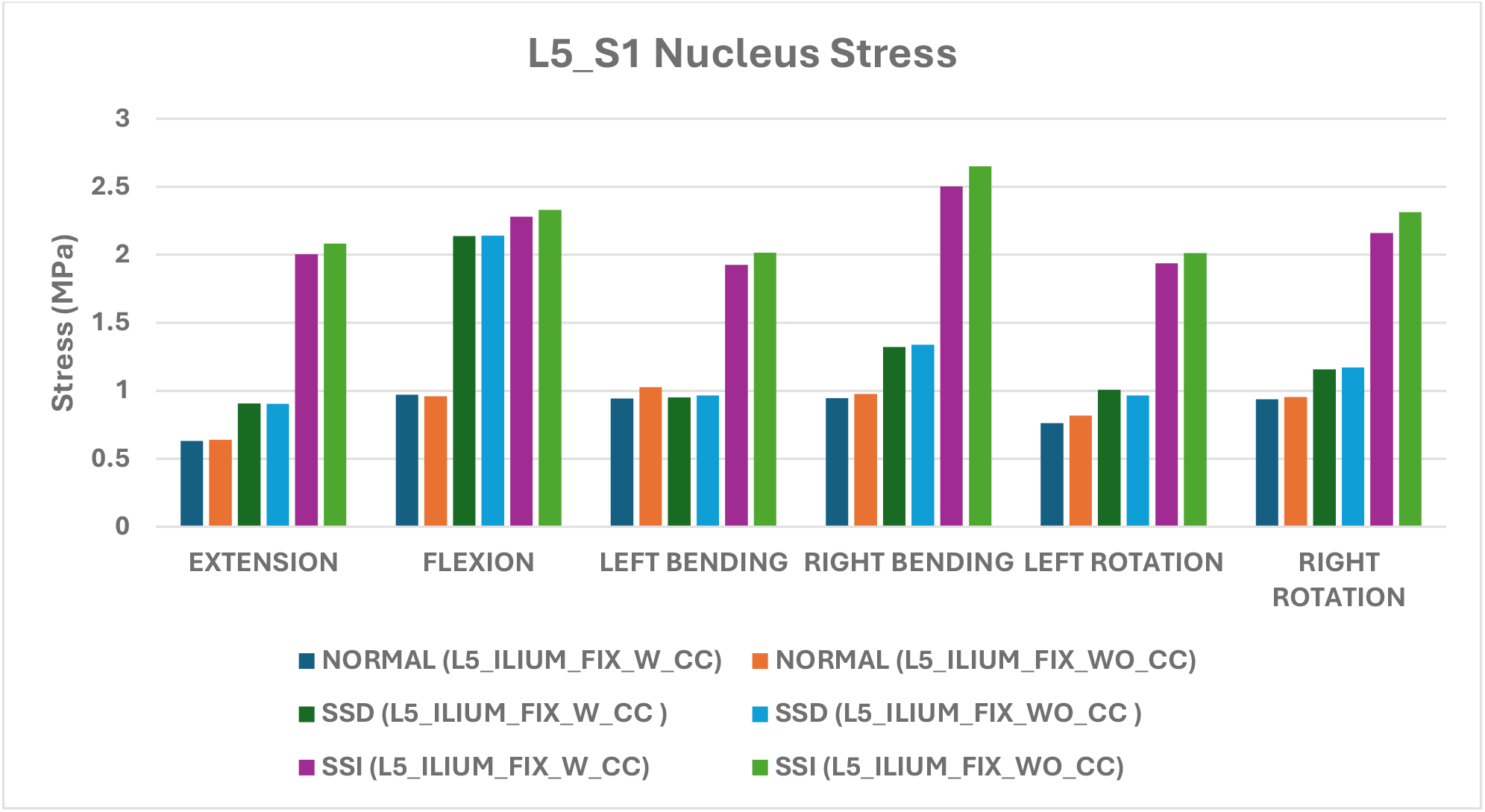
Comparison of L5-S1 Nucleus Stress at 7.5Nm moment with 400N follower load under two leg stance condition for three different sacral slopes stabilized with and without cross-connector.

#### Right SIJ Stress (Figure 8)

Models with higher sacral slopes showed higher SIJ stress (< 18 MPa) in both configurations for all loading conditions. Models with normal sacral slope demonstrated the least hip stress (< 12 MPa), followed by models with decreased sacral slope (<16 MPa) for both configurations and all loading conditions. Models with cross connectors demonstrated less SIJ stress than those without cross connectors for all loading conditions and cases.

**Figure 8.**
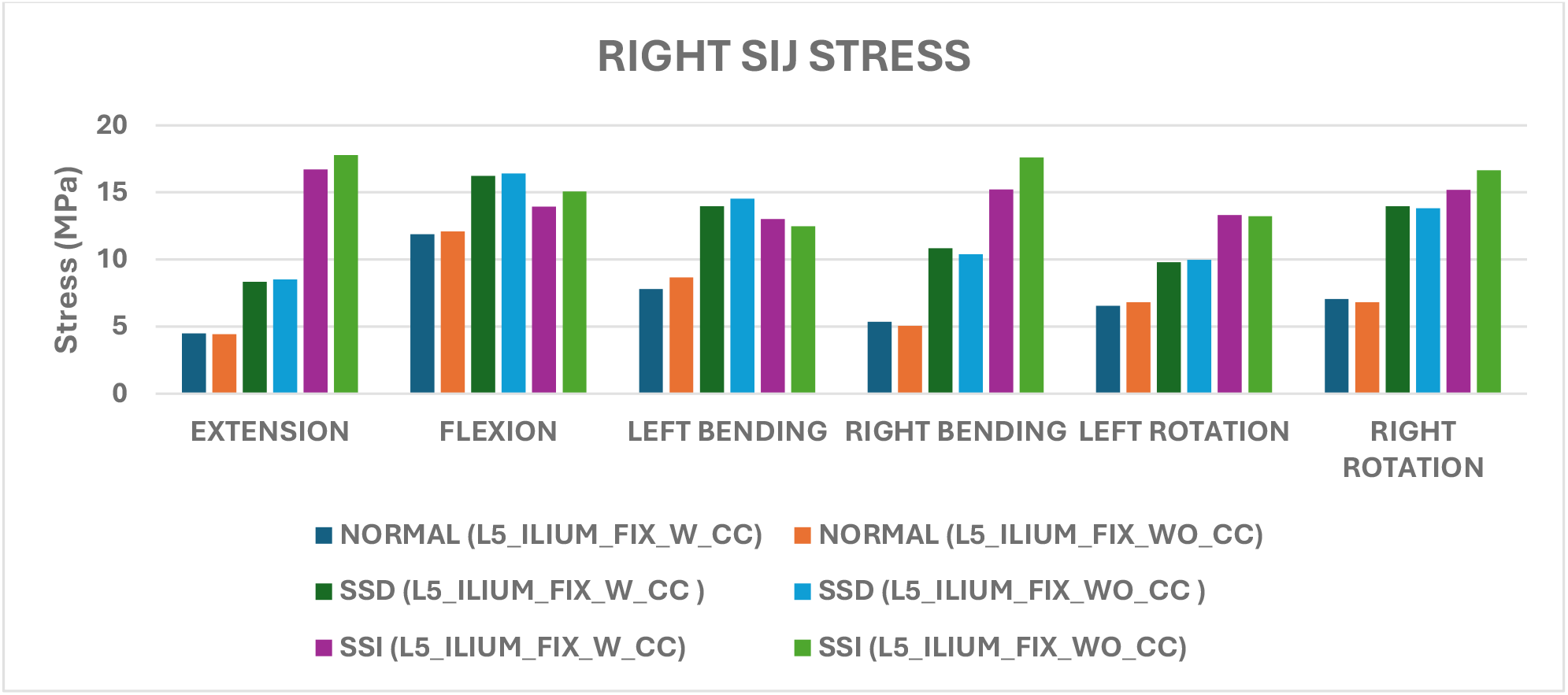
Comparison of Stresses at right SIJ at right SIJ at 7.5Nm moment with 400N follower load under two leg stance condition for three different sacral slopes stabilized with and without cross-connector.

### Displacement

#### Horizontal Displacement (Figure 9 and 10)

For the sacrum fracture, models with high sacral slopes recorded the least horizontal displacement (<0.6 mm), followed by decreased sacral slope models (< 0.5mm) for both configurations and loading conditions. Models with normal sacral slopes demonstrated the highest horizontal displacement (<1.4mm) for both configurations and all loading cases. Models with cross connectors recorded less horizontal displacement than models without cross connectors for both configurations and all loading conditions.

**Figure 9.**
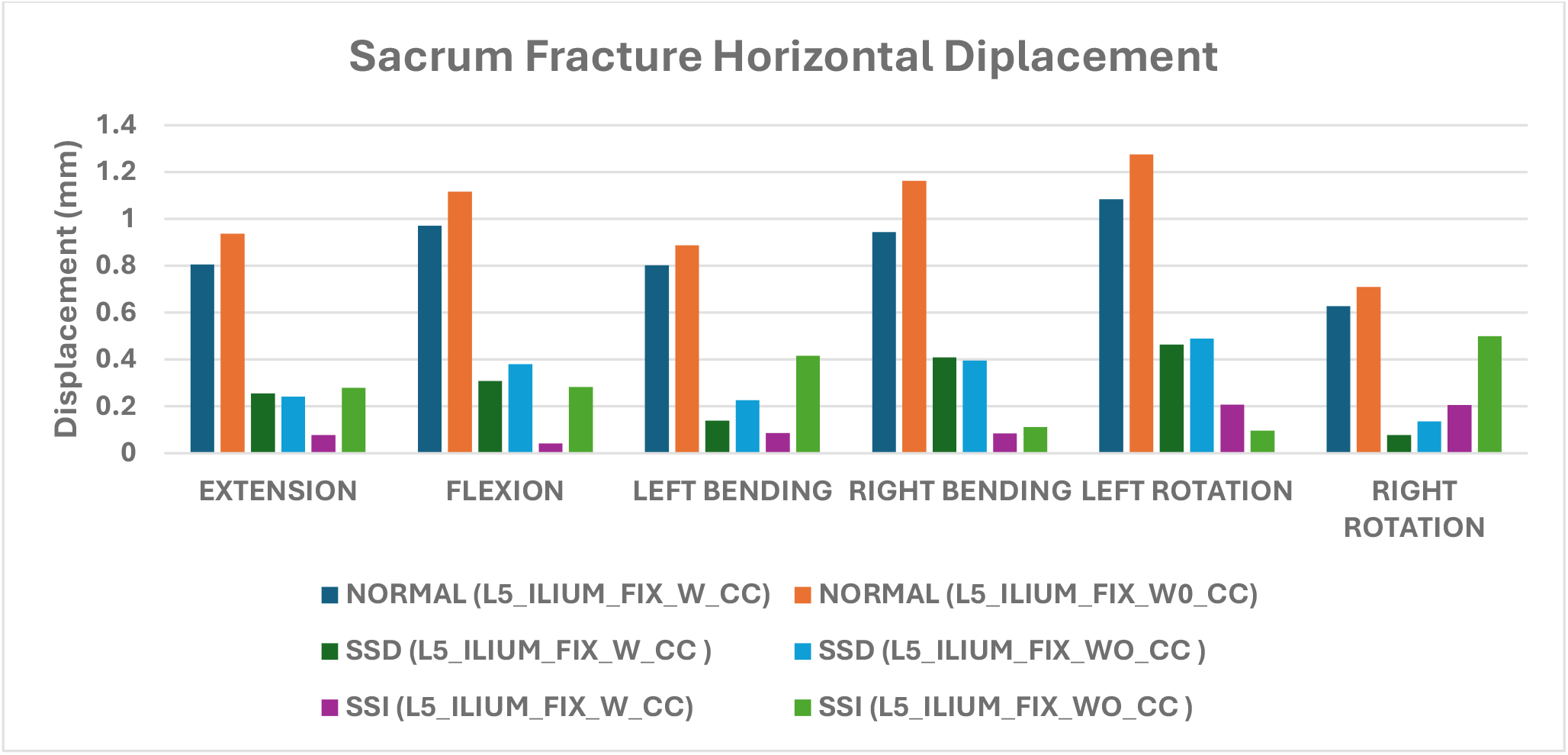
Comparison of Sacrum fracture horizontal displacement at 7.5Nm moment with 400N follower load under two leg stance condition for three different sacral slopes stabilized with and without cross-connector.

**Figure 10.**
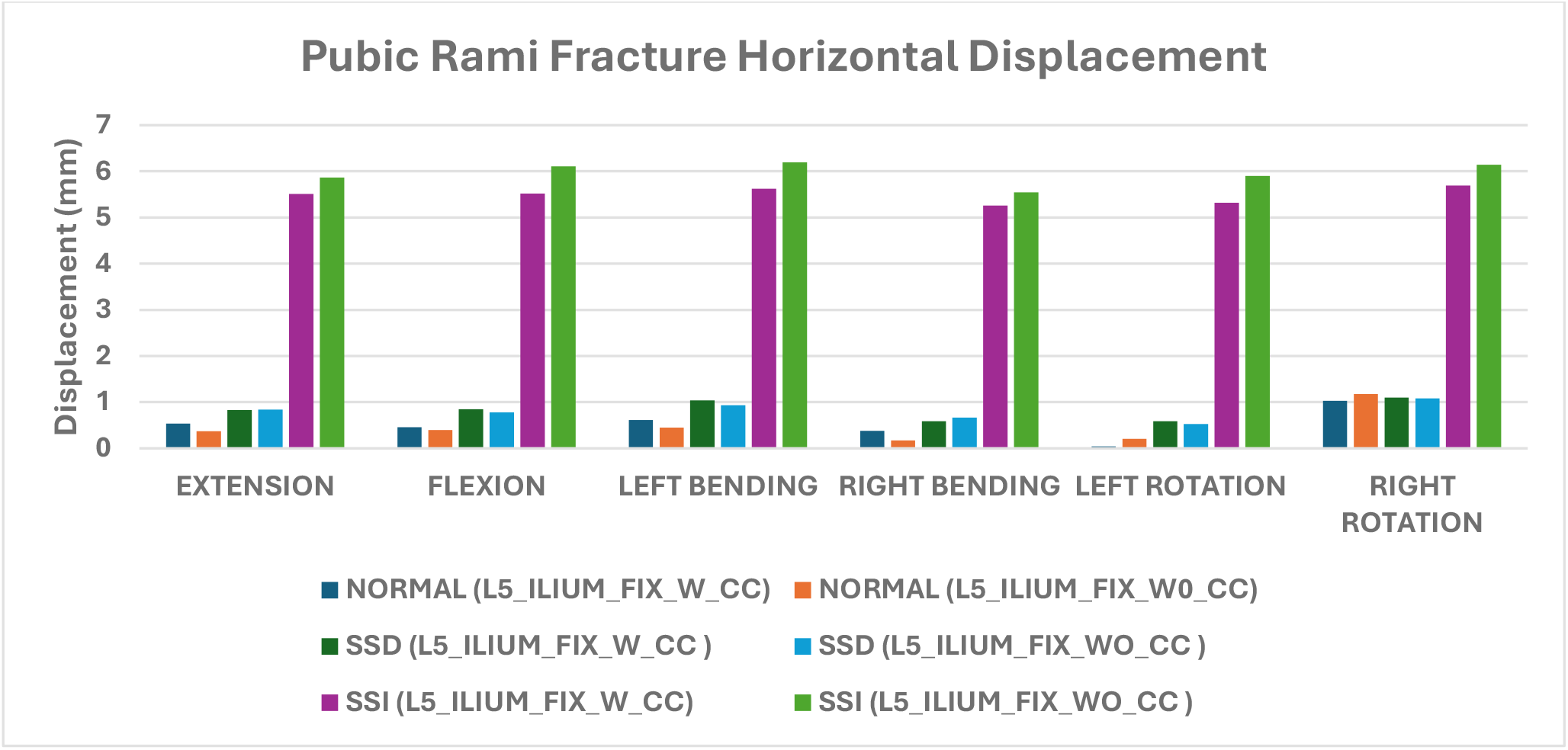
Comparison of Pubic Rami fracture horizontal displacement at 7.5Nm moment with 400N follower load under two leg stance condition for three different sacral slopes stabilized with and without cross-connector.

**Figure 11.**
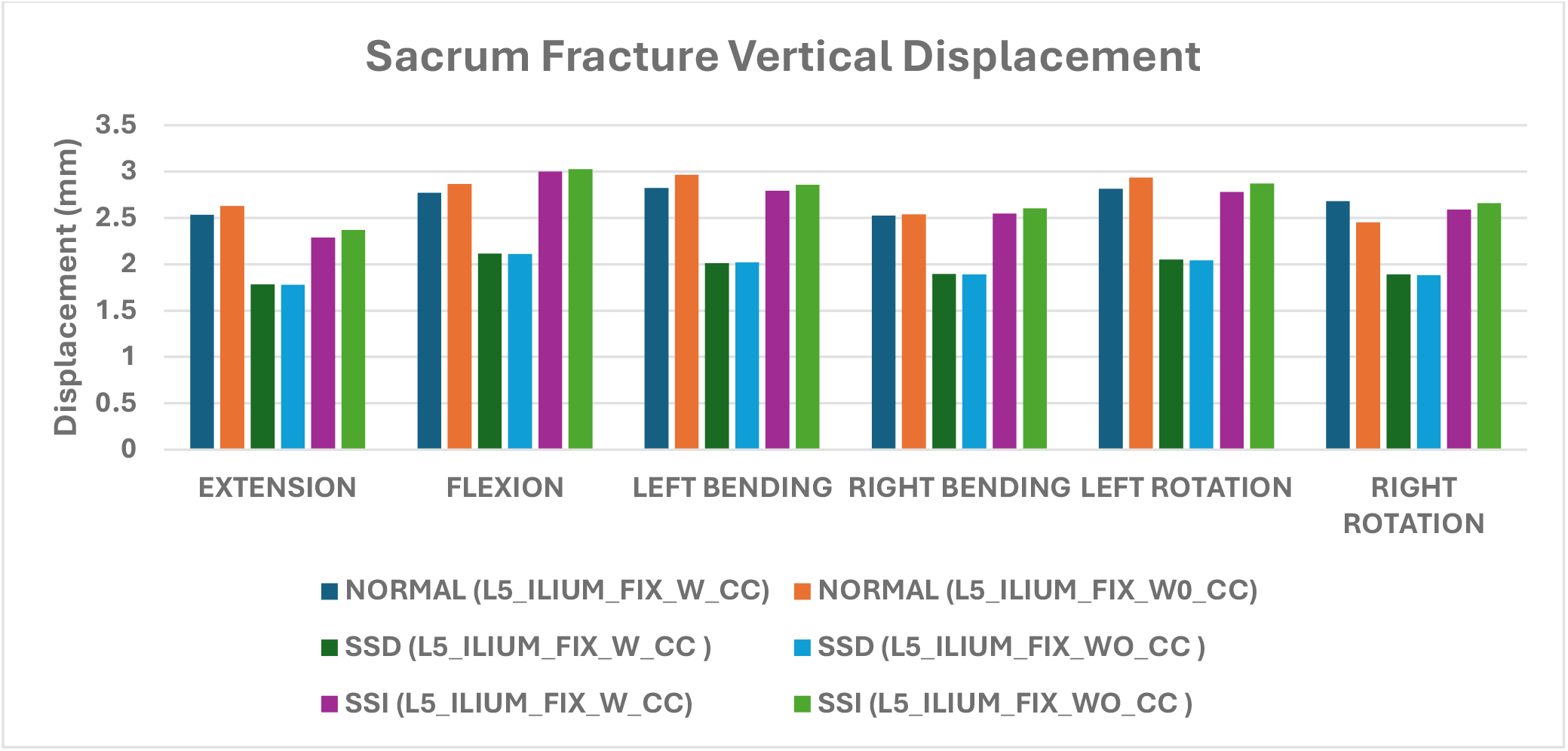
Comparison of Sacrum fracture vertical displacement at 7.5Nm moment with 400N follower load under two leg stance condition for three different sacral slopes stabilized with and without cross-connector.

For the pubic rami fracture, models with a normal sacral slope showed the least horizontal displacement (< 1mm), followed by the models with decreased sacral slopes (<2mm) for both the configurations and loading conditions. In contrast, models with higher sacral slopes showed a higher horizontal displacement (<7mm) for both configurations and all loading cases. Models with cross connectors recorded less horizontal displacement than models without cross connectors for both configurations and all loading conditions.

#### Vertical Displacement (Figure 11and 12)

Regarding the sacral fracture, models with a decreased sacral slope showed the least vertical displacement (<2mm), followed by models with a normal sacral slope (<2.5 mm) for all loading conditions and configurations. In comparison, models with a higher sacral slope measured the highest vertical displacement (<3mm) for all loading conditions and cases. Vertical displacement for models with and without cross connectors was similar for all the loading conditions and cases.

**Figure 12.**
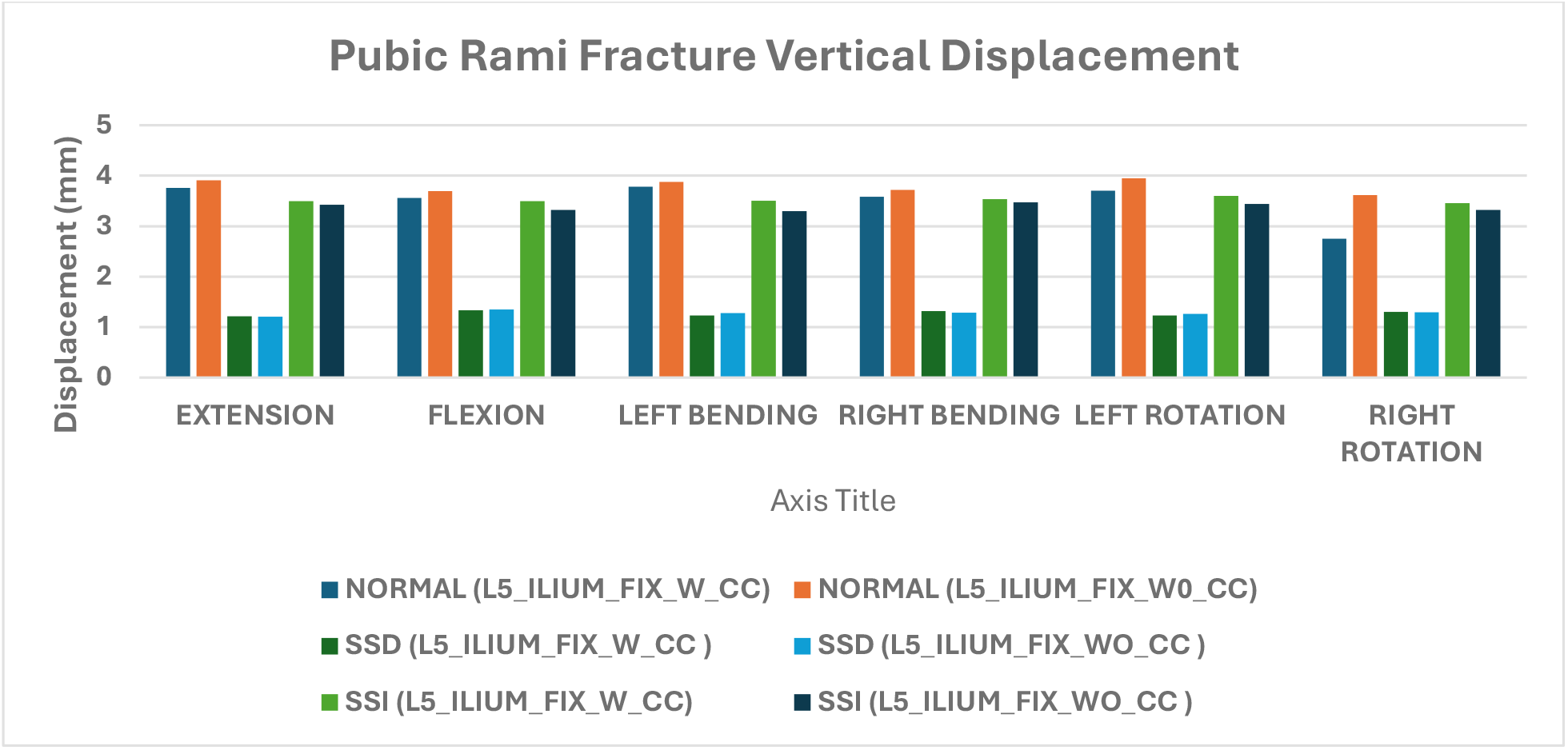
Comparison of Pubic Rami fracture vertical displacement at 7.5Nm moment with 400N follower load under two leg stance condition for three different sacral slopes stabilized with and without cross-connector.

Regarding the pubic rami fracture, models with decreased sacral slopes showed less vertical displacement (< 1.5mm). Vertical displacement for both the models with normal and increased sacral slopes recorded a similar displacement (< 4mm) for all the loading conditions and cases. Vertical displacement for models with and without cross connectors was similar for all the loading conditions and cases.

### Rod Stress (Table)

The model with decreased sacral slope treated with a cross connector showed the least rod stress (<200 MPa) for all loading conditions, followed by models with a normal sacral slope (<250 MPa) for both configurations for all loading conditions. Models with higher sacral slopes showed higher stress (< 350 MPa) for all loading conditions and cases. Models with cross connectors recorded least rod stress than models without cross connectors for both configurations and all loading conditions.

**Table 2.** Maximum von Mises Stress (MPa) on the implants for each motion simulated.

| Von Mises Stress (MPa) |  |  |  |  |  |  |
| --- | --- | --- | --- | --- | --- | --- |
|  | EXTENSION | FLEXION | LEFT BENDING | RIGHT BENDING | LEFT ROTATION | RIGHT ROTATION |
| NORMAL(L5_ILIUM_FIX_W_CC) | 210.8 | 243.8 | 262.8 | 184.7 | 232.2 | 210.3 |
| NORMAL(L5_ILIUM_FIX_WO_CC) | 217.4 | 304.1 | 268 | 242.4 | 238.6 | 215.1 |
| SSD (L5_ILIUM_FIX_W_CC) | 163.8 | 183 | 203.3 | 193.6 | 177.2 | 160 |
| SSD (L5_ILIUM_FIX_WO_CC) | 168.4 | 182.6 | 203.1 | 201.3 | 203.7 | 166.1 |
| SSI (L5_ILIUM_FIX_W_CC) | 298.5 | 341.3 | 312.2 | 335.5 | 333.7 | 309.7 |
| SSI (L5_ILIUM_FIX_WO_CC) | 290.2 | 326.6 | 304.7 | 340.9 | 287.2 | 298.2 |

## Discussion

The objective in this study was to investigate the biomechanical impact of pre-operative sagittal spinopelvic parameters, the SS and PT on the SIJ, hip joint and adjacent level IVD biomechanics after simulation of posterior fixation treatment for pelvic fracture. The effect of spino-pelvic alignment on posterior lumbopelvic fixation of pelvic fractures was analyzed with FE.

The rationale for selection of these posterior stabilization techniques was the previous study published by the author group whereby it was determined that these two posterior fixation techniques were superior in stabilizing sacral and contralateral sacroiliac joint range of motion as well as demonstrating stabilization in the horizontal displacement compared to vertical displacement of pubic rami fracture [25]. Several studies prior have performed biomechanical analyses to examine changes in the SS and its effect on instrumentation. Chen et al performed an FE analysis on posterolateral fusions (PLF) and posterior lumbar interbody fusion (PLIF) with SS angles of 35°-55°. They identified that patients with high SS angles who have isthmic spondylolisthesis may be better candidates for PLIF rather than PLF due to the high stresses seen on PLF instrumentation. These studies confirmed that increased SS negatively affect surgical instrumentation, increasing the stress on the sacral screws in PLF and PLIF models [26]. Similar trend was observed in this study where higher pre-operative sacral slopes led to increased stresses on the rods used in fixation of the simulated fractures.

Additionally, the models with increase SS showed highest amount of adjacent segment (L5-S1) ROM and IVD stresses. These trends are consistent with the author’s previous study with non-instrumented FE models as well as the observations from the studies conducted by Jiang et al, Berlemann et al, Drazin et al, and Vialle et al [27-30]. The studies have concluded that high SS patients may have increase risks of lumbar spondylolisthesis and disc degeneration at the lumbosacral level [13]. However, the addition of a cross connector led to the reduction of ROM and nucleus stress at the L5-S1 level when compared to lumbopelvic fixation without a cross connector. This may suggest that addition of cross connectors may aid in reducing the risk for adjacent segment degeneration following lumbopelvic fixation in patients with pre-existing high SS.

Regarding the SIJ biomechanics, published studies by Tonosu et al have showed that patients with high SS may be predisposed towards increased SIJ motion, and degeneration [31]. Our results showed that models with high SS showed the highest amount of SIJ motion on the contralateral side in flexion/extension and lateral bending. The addition of a cross connector to the posterior lumbopelvic construct showed the least amount of motion on the contralateral SIJ. The cross connector may help prevent loosening of the illiac screw on the contralateral side due to reduction in motion observed.

In this study, both the lumbopelvic fixation techniques in cases with normal SS showed most stability in terms of horizontal displacement at pubic rami fracture while the models with increased SS showed the highest amount of horizontal displacement at pubic rami fracture. Additionally, cases with decreased SS showed most stability in terms of vertical displacement at pubic rami fracture while the models with increase SS showed the highest amount of horizontal displacement at pubic rami fracture.

Conversely, when the horizontal displacements at sacral fractures were compared, the models with normal SS showed the highest amount of horizontal displacement at while cases with decreased SS showed most stability in terms of vertical displacement at sacral fracture. Vertical and horizontal displacements at the sacral fracture sites are correlated with significant longitudinal shear forces which in turn be associated with implant failures in previously published literature^33, 34^. The addition of a connector further increased the horizontal and vertical stability of the sacral fracture as well as at the public rami fracture site in all cases compared to the LP fixation without a cross connector.

When the stresses on the rods were compared, the models with increase SS showed the highest amount of rod stresses under all loading conditions. High stress regions on rods have been correlated with possible failure locations in previously published literature^32^. However, the addition of a cross connector reduced the rod stresses thereby reducing the possibility of instrument failures.

These results indicate patients with higher SS may be a risk factor for short term effects such as implant failures. Therefore, in such cases careful pre-planning needs to be done in case of choice of fixation when treating for unstable pelvic ring fractures. All lumbopelvic fracture fixation techniques in such cases may benefit from addition of cross connector due to increased facture site stability despite the risks from increased surgical time and surgical invasiveness Additionally, patients with higher SS may be at a long term risk for post-surgical degenerative effects such as L5-S1 degeneration, SIJ degeneration and hip joint degeneration. Therefore, pre-planning with regards to fracture fixation construct selection must be considered for such anatomic variations.

Lumbopelvic fixations have shown a potential for increase in adjacent segment degeneration per previously published clinical and biomechanical studies [32, 33]. Our results may suggest that lumbopelvic fixation with cross connectors may be beneficial for patients regardless of the pre-existing spinopelvic parameters, we suggest that these constructs should only be carefully selected in patients where other treatment options are not amenable [34].

### Limitations

This biomechanical study has some inherent limitations that need to be addressed. Firstly, complex muscle forces were simplified by substituting them with follower load on the spine, and linearly elastic material properties were assumed for implants and bones.A simplified version of Perfect fixation at the screw and bone interface as well as the screws, rods and cross connector interface was assumed without considering presence of any micromotions. The authors have assumed healthy bone properties in the simulated models. However, under osteogenic and osteoporotic bone properties the results presented in this study may not be applicable. The variation in spinopelvic parameters have been simplified to just 3 models and the authors are aware that more such combinations need to be simulated. Additionally, only posterior lumbopelvic fixation for treatment of unstable pelvic ring fractures have been modelled.

## Conclusion

The present study indicated that the change in SS affects the biomechanical stress of the lumbar spine, SIJ, hip joint, pelvic fracture, and instrumentations. Unstable pelvic fractures with large SS are more likely to cause instability at the fracture site and suggest that clinical studies should assess these FE outcomes following fixation.

